# Defining the Energetic Costs of Cellular Structures

**DOI:** 10.1101/666040

**Authors:** Gita Mahmoudabadi, Rob Phillips, Michael Lynch, Ron Milo

**Affiliations:** Stanford University; California Institute of Technology; Arizona State University; Weizmann Institute of Science

## Abstract

All cellular structures are assembled from molecular building blocks, and molecular building blocks incur energetic costs to the cell. In an energy-limited environment, the energetic cost of a cellular structure imposes a fitness cost and impacts a cell’s evolutionary trajectory. While the importance of energetic considerations was realized for decades, the distinction between direct energetic costs expended by the cell and potential energy that the cell diverts into cellular biomass components, which we define as the opportunity cost, was not explicitly made, leading to large differences in values for energetic costs of molecular building blocks used in the literature. We describe a framework that defines and separates various components relevant for estimating the energetic costs of molecular building blocks and the resulting cellular structures. This distinction among energetic costs is an essential step towards discussing the conversion of an energetic cost to a corresponding fitness cost.

---

Cells are out-of-equilibrium structures that require a constant supply of energy to remain in such a privileged state. Measuring the power required to run a cell or the heat produced as a cell goes through its normal metabolic operations is experimentally challenging (1-9). In addition to the challenges with measuring cellular power consumption, there are several plausible definitions for a cell’s rate of energy usage, making a rigorous discussion of the problem even more demanding. The interplay between the energetic cost of a cellular structure and its evolutionary fate is a subject that will become increasingly important as evolutionary cell biology matures as a science. Even though energetic censuses of cellular processes have been developed for decades (10-16), the distinction between direct energetic costs expended by the cell and the unoxidized substrates with energy production potential that the cell diverts into cellular biomass components, was not explicitly made. This has led to significant variation in the estimates for the energetic costs of molecular building blocks (2, 10, 12, 17-19).

Here, we clearly differentiate between various energetic costs incurred by a cell. The phrase “cellular structure” will be used as an umbrella term broadly referring to any cellular entity comprised of monomeric building blocks, be that entity a few base pairs of DNA, a protein complex, or a vesicle. At the heart of this subject are two important concepts: 1) all cellular structures have energetic costs composed of different components that should be properly defined and evaluated; and 2) the energetic costs and benefits of these cellular structures can translate into fitness differences that influence long-term evolutionary trajectories. We attempt to provide a quantitative framework for addressing the first issue and thus facilitate approaching the second issue. Indeed, the connection to organismal fitness (12, 17, 20) motivated the analysis described below, where we clarify the various components of energetic costs and thus note some of the important subtleties in performing energetic censuses of the cell.

Cellular energetic costs can be reported using several different units. We could, for example, report energetic costs in units of Joules. Under physiological conditions, ATP hydrolysis producing adenosine diphosphate (ADP) and orthophosphate (Pi) results in about −50 kJ/mol free energy change (21). However, the actual change in free energy depends on the exact concentrations of reactants and products. Because the hydrolysis of ATP and ATP-equivalent molecules serves as a universal currency of bioenergetics across the different domains of life (21, 22), and there is extensive biochemical knowledge regarding its investment in cellular processes, it is common to enumerate energetic costs in units of numbers of ATP hydrolyses (or their equivalent, e.g., the energetically similar GTP hydrolysis) (11, 18, 20, 23-25). To remain consistent, we will use the symbol P with different subscripts as a shorthand notation to represent an ATP (or an ATP-equivalent) hydrolysis event (10, 12, 25). Even with this seemingly straightforward approach, there still remain critical subtleties.

## The bioenergetic cost of a cellular structure

Cellular structures are assembled from molecular building blocks such as amino acids, nucleotides, lipids, and carbohydrates. If not provided by the outside environment, these monomeric subunits must be synthesized within the cell by processes requiring both carbon skeletons and the expenditure of energy. In fact, in many bacteria, all building blocks, as well as coenzymes and prosthetic groups can be synthesized from a small number of precursor metabolites (2). If some building blocks are available externally, the biosynthetic costs will be diminished, but there will still be costs of acquisition and transformation to arrive at the full set of internal building blocks (the cost of converting one amino acid to another, for example). In this paper, we will assume aerobic growth where all molecular building blocks are derived from a single carbon source, namely glucose, and further assume that sources of inorganic nitrogen and other trace elements are provided in excess within the growth media. These assumptions are especially applicable to growth conditions in the laboratory. We hope future studies will discuss costs for more complex media compositions. The assembly cost of a cellular structure must also include the requirements for construction of that structure from its molecular building blocks, e.g., the necessities for polymerizing a protein from its constituent amino acids, adding post-translational modifications, and folding the subsequent chain into the appropriate globular form. Finally, there will often be maintenance costs, e.g., maintaining the proton-motive force across the cell membrane, accommodation of molecular turnover, and identification and elimination of cumulative errors.

The sum of costs noted above represents the baseline investment that must be made in a cellular structure regardless of its benefit to the host cell (Figure 1). Given the near universality of many biosynthetic pathways and enzyme-reaction mechanisms, the assembly and maintenance costs can generally be calculated from information in the literature. The ability to make such calculations is a highly desirable complementary approach to laborious experimental approaches (26, 27), such as modifications of gene-expression levels, as these can have additional side effects (e.g., promiscuous binding or aggregation) that are difficult to quantify and irrelevant to construction/maintenance costs. What has been summarized in the paragraphs above, however, are the **direct costs** of a cellular structure, which do not fully describe the energetic consequences for the cell. We use PD to symbolize this direct cost (Figure 2).

**Figure 1.**
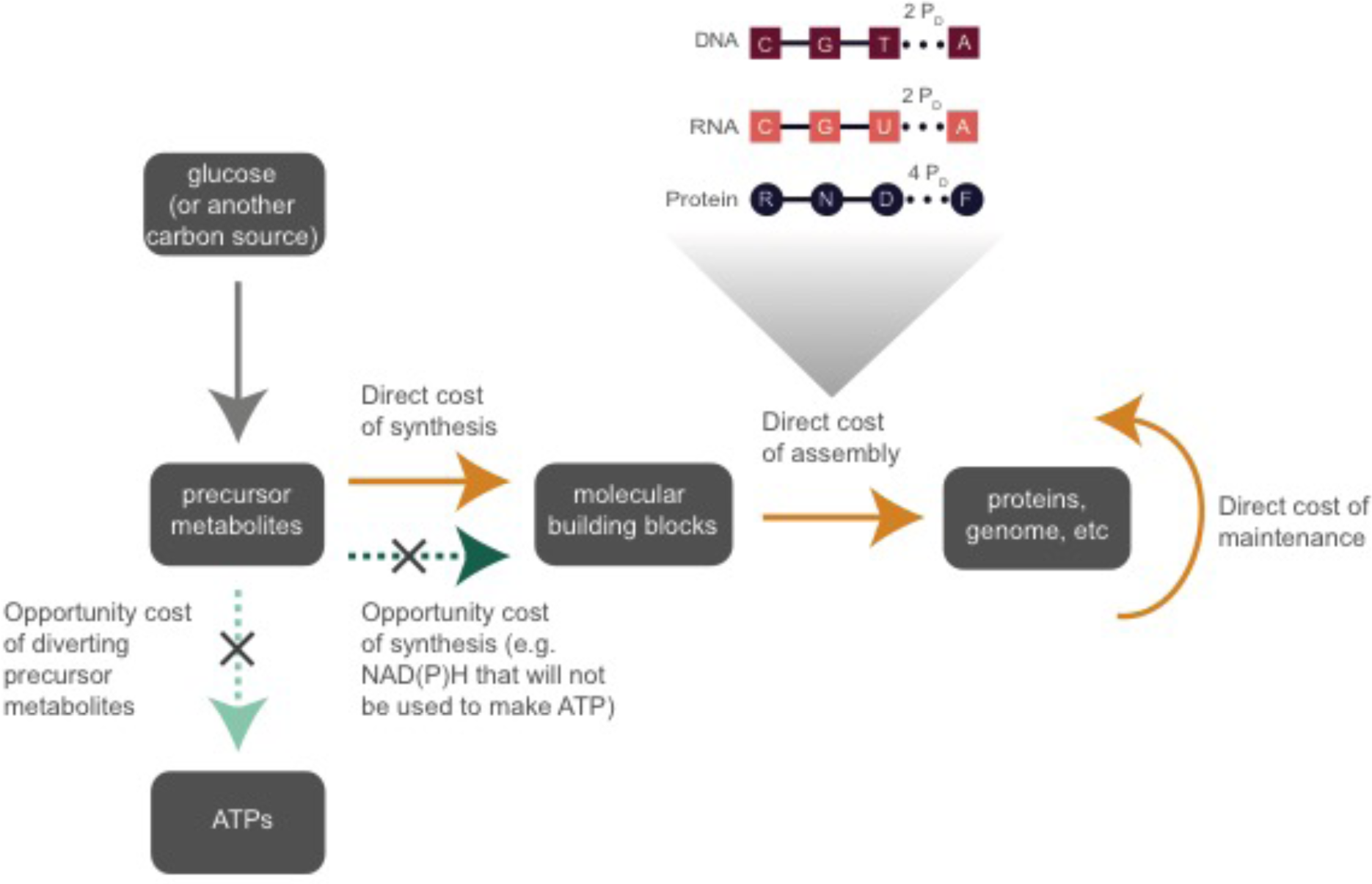
The distinction between direct and opportunity costs associated with synthesizing molecular building blocks. As glucose is partially metabolized into precursor metabolites, the energy that could have been captured from the complete metabolism of glucose is referred to as the opportunity cost of precursor metabolites (denoted by a light green arrow). Conversion of precursor metabolites into molecular building blocks consumes electron carrier molecules such as NADH, which are not counted in the direct cost as there is no actual hydrolysis reaction that releases heat. If not used in synthesis of building blocks, these electron carriers would have generated ATP, and thus the conversion of precursor metabolites to building blocks incurs additional opportunity cost (giving rise to the second, darker, arrow). The conversion also consumes ATP, which we count as the direct cost of synthesis. The assembly of macromolecules such as proteins from building blocks requires additional post-synthesis costs such as the cost of polymerization and maintenance. The polymerization costs per nucleotide or amino acid are denoted. Direct costs are shown by solid orange arrows, whereas the opportunity costs are denoted in shades of green and dotted lines. The same color scheme is used in Figure 2.

**Figure 2.**
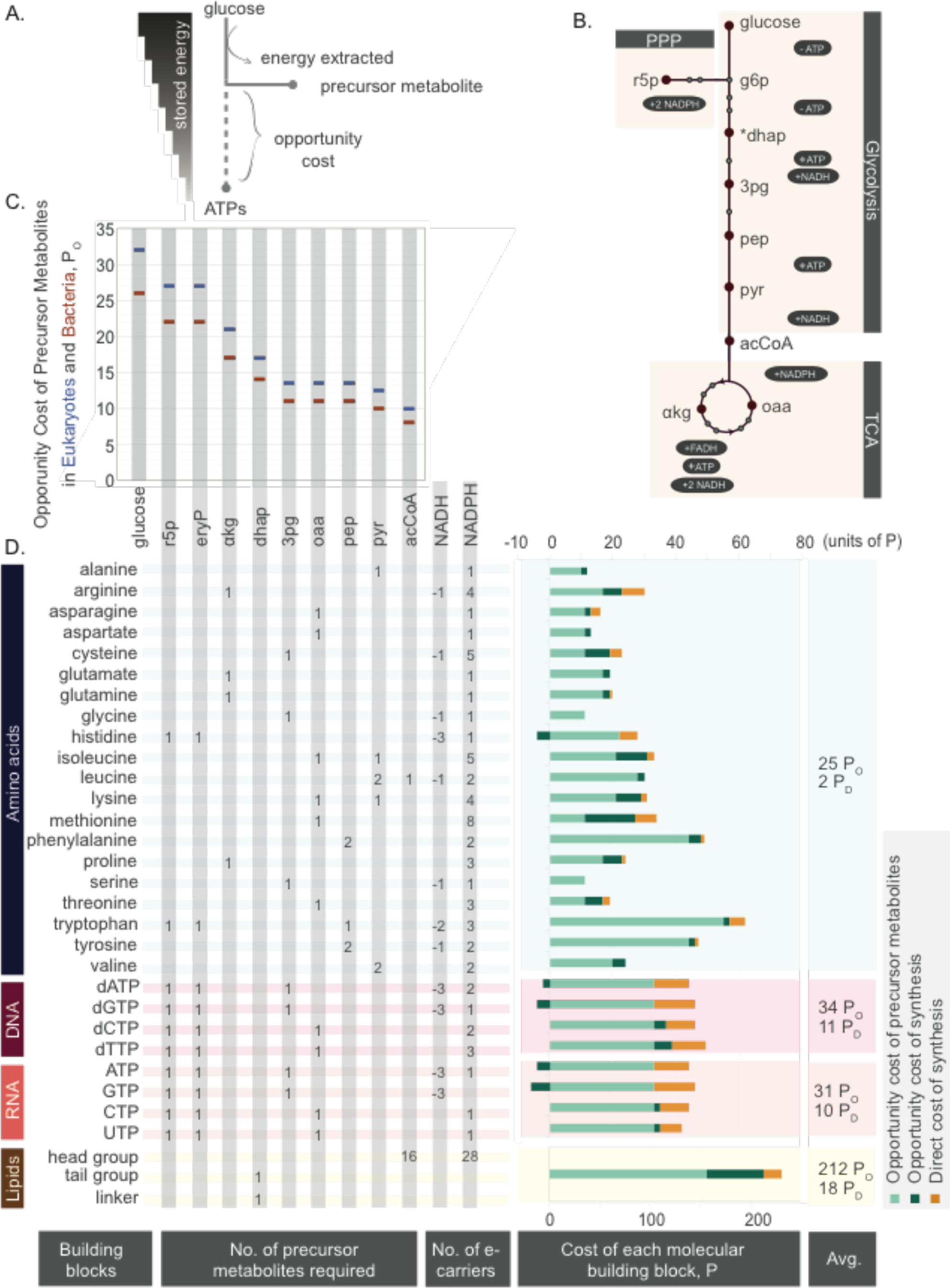
The cost of molecular building blocks. **A)** The concept of opportunity cost is shown schematically for the situation in which glucose is the sole carbon source. **B)** The metabolic pathways from which the opportunity cost of each precursor metabolite can be estimated. From dihydroxyacetone phosphate (dhap) onwards, there are two molecules of each precursor metabolite generated. Precursor metabolites that are not implicated in the synthesis of building blocks are shown as grey circles. Positive and negative signs indicate gains and losses in ATPs or electron carrier molecules from the conversion of one precursor metabolite to another. The names for each precursor metabolite denoted are as follows: ribose-5-phosphate (r5p), erythrose-4-phosphate (e4p, or eryP), alpha-ketoglutarate (akg), dihydroxyacetone phosphate (dhap), 3-phosphoglycerate (3pg), oxaloacetate (oaa), phosphoenolpyruvate (pep), pyruvate (pyr), acetyl-CoA (acCoA). **C)** The opportunity cost of each precursor metabolite estimated in the context of both heterotrophic bacteria and eukaryotic cells as detailed in the SI. **D)** The opportunity cost of a molecular building block is equivalent to the number of precursor metabolites and electron carrier molecules used during its synthesis times their respective opportunity costs. The direct cost of synthesizing a molecular building block is the number of ATP (or ATP-equivalent) hydrolysis events required during the synthesis of each building block (orange). All costs shown are estimated in the context of a heterotrophic bacterial metabolism. The average direct cost provided for building blocks does not include post-synthesis costs such as polymerization or maintenance. For lipids we use a representative bilayer building block out of the natural diversity. See SI Table 1 for further details.

The construction and maintenance of cellular structures represent a drain on resources that could otherwise be allocated to other cellular functions. As metabolic precursors that can be fully metabolized for ATP production are instead allocated as carbon skeletons to the production/maintenance of a particular trait, this diversion eliminates their availability for other purposes, a consequence that we refer to as the **opportunity cost**. We use PO to symbolize this opportunity cost (Figure 2).

Opportunity costs can also be calculated from basic cell-biological knowledge (10, 17, 18), at least for simple cases such as growth on glucose. Specifically, we estimate the opportunity cost of a precursor metabolite as the number of ATPs (or ATP equivalents) that could have been generated had the precursor metabolite not been diverted towards the synthesis of molecular building blocks (Figure 2A, SI). Figure 2B demonstrates the placement of metabolic precursors that are important in the synthesis of molecular building blocks across metabolic pathways. In arriving at cost estimates for molecular building blocks, we assumed glucose as the primary carbon source under aerobic conditions. These cost estimates will need to be modified when considering another carbon source as input. The estimated opportunity costs for each precursor metabolite in the context of bacterial and eukaryotic metabolism are shown in Figure 2C. The difference in these costs between eukaryotic and bacterial metabolism stems from the higher efficiency of eukaryotic metabolism in producing ≈6 more ATPs per glucose than bacterial metabolism, which generates about 26 ATPs per glucose (28, 29). We hope that future studies will also reveal these costs in the context of archaeal metabolism.

Another source of opportunity cost is the pool of electron carrier molecules used in the synthesis of molecular building blocks. If electron carriers such as NAD(P)H were to be preserved rather than used in the synthesis of molecular building blocks, they would each result in about 2 ATP molecules within the bacterial electron transport chain (or 2.5 in eukaryotic systems, see SI) (29). To distinguish between these two sources of opportunity cost, we have denoted them in two shades of green in Figures 1 and 2. However, unless denoted otherwise, we will refer to the opportunity cost of a molecular building block as the sum of all opportunity costs.

The **total cost** of producing a trait and diverting structures from alternative usage to do so is the sum of the direct and opportunity costs,

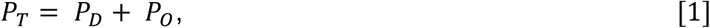

where all costs represent the cumulative expenditures over the entire lifespan of the cell. Whereas *P*_*D*_ is expected to reflect ATP (and ATP-equivalent) hydrolysis reactions resulting in heat dissipation in the cell, *P*_*O*_ will not be manifested in heat production, given that the ATP is not actually produced or consumed. We have used the symbol PT to denote the number of ATP formation/hydrolysis reactions associated with the total cost definition (Figure 2).

### Which is more biologically relevant, the direct or the total cost definition?

Depending on the experimental context or question at hand, one of the two definitions can be more appropriate than the other. Under the direct-cost definition, we simply account for the number of ATP hydrolysis events, and ignore the effects of diverting molecular building blocks from energy-producing pathways. The direct cost is also of interest in scenarios where there is a constrained rate of ATP production, e.g., because of a limited amount of membrane real-estate for the respiration machinery (30) or a limitation on the amount of glycolytic enzymes under anaerobic conditions. Most of the pioneering studies in bioenergetics have reported the costs of molecular building blocks in terms of their direct costs (2). More recent works on cellular energetics, however, have implicitly adhered to the total cost definition, thereby including the opportunity costs of molecular building blocks (10, 12, 17-19).

The total cost definition may be useful when the overall carbon source availability is limiting, under which the implication of diverting a substrate for biomass buildup, instead of using it for ATP production, could affect fitness just as well as a need to invest ATP. Such a situation is encountered in the lab during growth in carbon-limited chemostats, which is also found to be akin to many natural habitats (31). Growth in chemostats has the advantage of enabling precise carbon accounting. During a chemostat experiment, the number of glucose molecules required per unit time to grow cells at a set growth rate can be measured. This number can then be converted to an energetic value by assuming that every glucose molecule consumed by the cell is fully metabolized. However, not all glucose molecules are used for energy production. In fact, as we will briefly discuss in the following section, depending on the growth rate, the majority of the carbon atoms in the glucose molecules consumed will be converted to biomass components rather than respired to CO_2_. As a result, the cost of a cell as measured through substrate usage accounting (as performed in many chemostat experiments) includes both the direct and the opportunity costs of cellular processes, and is inherently a total cost estimate. Thus, when considering the energetic burden of a given process on a cell’s energy budget as measured by chemostat experiments, the total cost definition provides a more meaningful approach.

### What fraction of a cell’s total cost is direct cost?

Knowledge of the dry mass of a cell gives an estimate of the mass of carbon diverted to biomass production instead of ATP production and can thus be useful in a rough estimate of the total cost, PT. An *E. coli* cell with an approximate volume of 1 µm^3^ (32) and ≈50% carbon in its dry weight contains ≈0.1 pg carbon (5 × 10^9^ carbon molecules). Growing on glucose as its sole carbon source and considering that every glucose molecule contributes 6 carbons, we can deduce that the demands for carbon of this bacterium (for biomass skeletons) is met by ≈10^9^ glucose molecules. Because every glucose molecule can generate about 26 ATPs during aerobic respiration in *E. coli*, we can convert the number of glucose molecules thus used for the biomass skeleton to an opportunity cost of ≈3 × 10^10^ PO.

To obtain the direct cost estimate of a bacterial cell, we can make use of the fact that translation incurs the greatest cost for cells (10-12, 17) (this situation also holds true for viral infections (25, 33)). An *E. coli* cell with a volume of 1 µm^3^ contains about three million proteins (34). With an average protein length of 300 amino acids (35), an *E. coli* cell of this size will be comprised of 10^9^ amino acids. The direct costs of polymerization (4 PD) and synthesis from precursor metabolites (≈2 PD) (Figure 2) are ≈6 ATPs per amino acid, and thus the direct cost of translation is about ≈6 × 10^9^ PD (for comparison, the direct cost of genome replication for an *E. coli* genome comprised of 5 × 10^6^ base pairs is ≈10^8^ PD); this is because the direct cost of polymerization (2 PD) and synthesis of nucleotides from precursor metabolites (11 PD) amounts to 13 PD per nucleotide). Taking the direct cost of translation as a proxy for the sum of all direct costs, the ratio of direct cost of an *E. coli* cell (≈6 × 10^9^ PD) to its opportunity cost (≈3 × 10^10^ PT) is 0.2. This suggests that the majority of glucose molecules consumed by an *E. coli* cell during rapid growth are not fully metabolized and instead are used to synthesize biomass. Of course, this is a very crude analysis that does not differentiate between the costs of nucleotides, lipids, proteins, or the cell wall. Yet the reasoning may be useful, and the dominance of proteins in the biomass composition makes the conclusion robust. Moreover, the values derived in this estimate for the cost of *E. coli* are similar to those obtained from growth experiments in chemostats (12).

## Energy as one of several possible limiting factors to growth

Cellular fitness need not always be strongly limited by energy. For example, plants and photosynthetic plankton populations often experience an overabundance of energy availability relative to nutrients such as nitrogen, phosphorus or iron. For an organism living in an environment plentiful in a carbon and energy source but limited by another nutrient, energy extracted from the food source may be in excess supply, resulting in under-utilization of ingested carbon and energy. The energetic accounting in the usage of the carbon substrate can still be evaluated as given here in terms of the direct and opportunity costs but their effect on growth will be diminished due to the greater impact of the other nutrient shortage on growth rate.

## Conclusion

Thanks to a wealth of classical experiments in biochemistry from several decades ago, we have access to detailed information about metabolic pathways. Despite this vast body of knowledge, the interplay of energetics and evolution has largely remained unexplored. In this paper, we attempt to clarify the existence of different components and definitions for the energetic cost of cellular structures. Our work could help dedicated future efforts of translating the energetic cost of a cellular structure to a fitness cost which would enable quantitative predictions about the evolutionary trajectories of cellular structures in energy-limited conditions.

## Acknowledgement

We are grateful to Michael Lassig for his comments and insightful recommendations. G.M. is supported by the National Science Foundation Graduate Research Fellowship (grant no. DGE-1144469). R.M. is supported by the European Research Council (Project NOVCARBFIX 646827), the Israel Science Foundation (grant No. 740/16), and the Beck-Canadian Center for Alternative Energy Research. R.P. is supported by The John Templeton Foundation (Boundaries of Life Initiative, grant ID 51250), the National Institute of Health’s Maximizing Investigator’s Research Award (grant no. RFA-GM-17-002), the National Institute of Health’s Exceptional Unconventional Research Enabling Knowledge Acceleration (grant no. R01-GM098465), and the National Science Foundation (grant no. NSF PHY11-25915) through the 2015 Cellular Evolution course at the Kavli Institute for Theoretical Physics. M.L. is supported by the U.S. Department of Army, MURI award (grant no. W911NF-14-1-0411), the National Institutes of Health (grant no. R35-GM122566) and the National Science Foundation (grant no. MCB-1518060).

## Supplementary Information

**SI Table 1**. A detailed breakdown of the energetic cost structures depicted in Figure 2.

### Opportunity Costs of Precursor Metabolites

In this section we will continue to use the symbol P to refer to ATP or ATP-equivalent hydrolysis events. Once we arrive at the opportunity cost of each precursor metabolite, we will denote these costs by the symbol P_O_ in Figures and main text to clearly distinguish from any direct costs. The derivations we used for the opportunity cost of precursor metabolites in heterotrophic bacteria versus eukaryotes is very similar, with the following difference: the total number of ATPs generated per glucose during aerobic respiration, *E*_*glu*_ are 26 P (28) in bacteria and 32 P in eukaryotes. This is because, from each molecule of NAD(P)H and FADH_2_, the mitochondrial electron transport chain produces 2.5 P and 1.5 P, whereas the bacterial electron transport chain produces 2 P and 1 P, respectively (2, 29, 36). It is important to note that the theoretical yield often mentioned of 38 P per glucose for eukaryotes is based on the assumption that each NAD(P)H and FADH_2_ results in 3 P and 2 P (29) which has been shown to be inaccurate. Here, we will carry out estimates with *E. coli* in mind, however these calculations can be similarly carried out for heterotrophic eukaryotes (Figure 2.C) (25).

To arrive at the opportunity cost of precursor metabolites, we have to consider the partial gain or loss of energy that results from the conversion of glucose to a precursor metabolite. We will refer to the number of ATP molecules generated in this process as *E*_*partial*_ (Figure 2.A). We further define the opportunity cost of a precursor metabolite, *C*_*opportunity*_, as

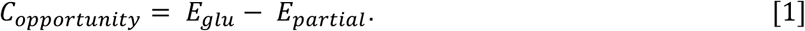

*C*_*opportunity*_ will depend on where in a metabolic pathway a precursor metabolite is generated. In most cases, the conversion of glucose into a precursor metabolite will result in a net energy gain, but in some cases the conversion will induce a net energy loss. A good example is dihydroxyacetone-phosphate (dhap), a precursor metabolite required during lipid synthesis (Figure 2.B). Each glucose molecule results in the production of two dhap molecules, with this process having a net energetic cost of 2 P (*E*_*partial*_= -*2* P) (Figure 2.B). Thus, based on SI Eq.1 we can conclude that the opportunity cost of two dhap molecules is 2 P greater than that of glucose (28 P), or 14 P per dhap molecule (Figure 2.C). We will now carry out similar calculations for other precursor metabolites using the information about metabolic pathways shown in Figure 2.B. We have provided a summary of these derivations in Figure 2.C.

To simplify SI Eq. 1, we could simply account for the change in energy as we convert one precursor metabolite into another. Specifically, we can derive the opportunity cost of metabolite *j*, 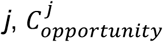 from the opportunity cost of metabolite *i*, 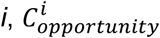 by

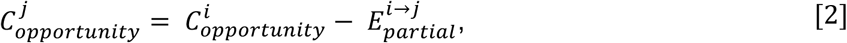

where 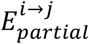 corresponds to the net energy gain resulting from the conversion of metabolite *i* to metabolite *j*. During glycolysis each molecule of dhap is converted to a molecule of 3-phosphoglycerate (3pg), resulting in the production of 1 NADH molecule and 1 ATP. Because each NADH is equivalent to 2 P, the net energetic gain from this conversion is 3 P. Therefore, the 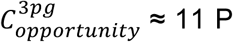 (SI Eq. 2) (SI Figure 2B). If not used as a precursor, 3pg is converted to phosphoenolpyruvate (pep), in a process that has a net energy gain of zero. Therefore, 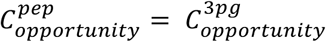. In converting pep into pyruvate, there is a net energy gain of 1 P, so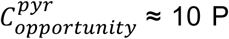 Pyruvate can be converted to oxaloacetate (oaa) with the expenditure of 1 P, hence, 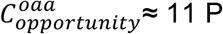

Pyruvate is further converted to acetyl-CoA (acCoA), and in the process one molecule of NADH is generated. The 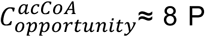 because each NADH is equivalent to 2 P in bacteria. One molecule of acCoA and one molecule of oaa are then eventually converted to alpha-ketoglutarate (akg). The sum of the opportunity costs of acCoA and oaa is 19 P, and because 1 molecule of NADH is generated in their conversion to αkg,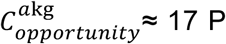 Similarly, αkg is eventually converted to oaa, and 2 NADH, 1 GTP, and 1 FADH_2_ molecules are generated. With a net energy gain of ≈ 6 P from this conversion,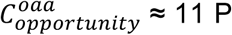, which is consistent with the opportunity cost of oxaloacetate derived from the anaplerotic pathway described earlier (the conversion of pyruvate to oxaloacetate via the pyruvate decarboxylase enzyme).

Glucose can also be converted to ribose-5-phosphate (r5p) in the pentose phosphate pathway, and in the process 2 NADPH molecules are generated, which is equivalent to 4 P. We subtract 4 P from the possible 26 P that glucose would be converted to under respiratory conditions, and we arrive at 22 P as the opportunity cost of ribose-5-phosphate. The same calculation can be used for erythrose-4-phosphate (e4p), resulting in 22 P as its opportunity cost.

